# Urine proteomic analysis of the rat e-cigarette model

**DOI:** 10.1101/2022.11.19.517186

**Authors:** Yuqing Liu, Ziyun Shen, Chenyang Zhao, Youhe Gao

**Affiliations:** Gene Engineering Drug and Biotechnology Beijing Key Laboratory, College of Life Sciences, Beijing Normal University, Beijing 100875, China

**Keywords:** urine, proteomics, e-cigarette model, odorant-binding protein

## Abstract

Urinary proteomics was used to investigate the potential effects of e-cigarettes on the human body. In this study, a rat e-cigarette model was constructed by smoking for two weeks and urine samples before, during, and after e-cigarette smoking were collected. Urine proteomes before-after smoking of each rat were compared individually, while the control group was set up to rule out differences caused by rat growth and development. After smoking, the differential proteins produced by rats shows strong individual variation. Fetuin-B, a biomarker of COPD, and annexin A2, which is recognized as a multiple tumor marker, were identified as the differential proteins in five out of six smoking rats on day 3. To our surprise, odorant-binding proteins expressed in the olfactory epithelium were also found and were significantly upregulated, which may help explain olfactory adaptation. Pathways enriched by the differential proteins shows the evidence that smoking e-cigarettes affects the immune system, cardiovascular system, respiratory system, etc., which provides clues for further exploration of the mechanism of e-cigarettes on the human body.

## 1. Introduction

### 1.1 E-cigarettes

E-cigarettes are mainly composed of four parts: soot oil, heating system, power supply and filter nozzle. Aerosols with specific odors are generated by heating atomization for smokers. The main components of aerosol liquid of electronic vaporizer are plant glycerin, propylene glycol, edible flavor and nicotine salt. As of 2019, approximately 10 million people aged 15 years and older in China have used e-cigarettes^[1]^. The population using e-cigarettes is predominantly young adults, with the highest use in the 15- to 24-year age group^[1]^. The vast majority (58.3%) of middle school e-cigarette users use fruit-flavored e-cigarettes, while previous research suggests that these tastes may attract young people to try e-cigarettes^[2]^. On May 1, 2022, *the Regulations on the Administration of Electronic Tobacco* prohibited the use of electronic cigarettes other than tobacco tastes. Pipe AL et al.^[3]^ found that the composition of heated chemical aerosols inhaled by the human body after electron fumigation is very complex, including nicotine, nitrosamines, carbonyl compounds, heavy metals, free radicals, reactive oxygen species, particulate matter, and “emerging chemicals of concern”, which further demonstrates the potential harm of smoking e-cigarettes. Studies have shown that smoking e-cigarettes may increase the risk of lung disease^[4]^ and cardiovascular disease^[5][6]^ and may cause harm to the liver^[7]^, urinary system^[8]^, and immune system^[9]^. Smoking e-cigarettes not only causes harm to themselves but may also cause harm to the foetus in pregnant women exposed to e-cigarettes. BallbèM et al.^[10]^ detected low but nonnegligible concentrations of e-smoke-associated analytes in cord blood and breast milk of nonuser pregnant women exposed to e-cigarettes. Aslaner DM et al.^[11]^ also demonstrated that the inhalation of second-hand e-cigarette smoke by pregnant women can have long-term effects on the lungs of offspring. At the same time, because the nicotine content in e-cigarette smoke is equivalent to, or even higher than, that in the combustible smoke^[12]^, the drug addiction damage caused by e-cigarette smoke to the human body cannot be ignored.

### 1.2 Urine Biomarkers

Biomarkers are indicators that can objectively reflect normal pathological processes as well as physiological processes^[13]^, and clinically, biomarkers can predict, monitor, and diagnose multifactorial diseases at different stages^[14]^. The potential of urinary biomarkers has not been fully explored compared to more widely used blood biomarkers, especially in terms of early diagnosis of disease and prediction of status. Because homeostatic mechanisms are regulated in the blood, changes in the blood proteome caused by disease are metabolically excreted, and no significant changes can be apparent in the early stages of the disease. Whereas urine is produced by glomerular filtration of plasma and is not regulated by homeostatic mechanisms; thus, minor changes in the disease at an early stage can be observed in urine^[15]^, which shows that urine is a good source of biomarkers.

Currently, the detection of biomarkers in urine has attracted increasing attention from examiners and researchers. This approach has been used in the treatment and research of a variety of diseases such as pulmonary fibrosis^[16]^, colitis^[17]^, glioma^[18]^ and other diseases. Studies have shown that urine biomarkers can classify diseases such as predicting chronic kidney disease (CDK)^[19]^ and distinguishing benign and malignant ovarian cancer^[20]^. Urine biomarkers can also be used to detect whether complete resection and recurrence occur after tumor surgery so that adjustments can be made in time to reduce the risk of recurrence. Pharmacologically, urinary biomarkers can observe the utility of drugs on the body, such as predicting the efficacy of rituximab therapy in adult patients with systemic lupus erythematosus (SLE), and sacubitril-valsartan is more effective than valsartan in the treatment of chronic heart failure^[21]^. In terms of exercise physiology, urine biomarkers can reflect changes in urine proteomics after exercise, thus providing a scientific basis for the rational training of athletes^[22]^. In recent years, many studies have shown that urine proteomics can also reveal biomarkers in psychiatric diseases, such as parkinsonism^[23][24]^, Alzheimer’s disease^[25]^, depression^[26]^, autism^[27]^ and other diseases.

However, there have been no studies on e-cigarettes in the field of urine proteomics. The urine proteome is susceptible to multiple factors, such as diet, drug therapy, and daily activities. To make the experimental results more accurate, it is critical to use a simple and controllable system. Because the genetic and environmental factors of animal models can be artificially controlled and can minimize the influence of unrelated factors, the use of animal models is a very appropriate experimental method. Therefore, we constructed an animal model to analyze the urine proteomics of the rat e-cigarette model, and the experimental workflow is shown in Fig. 1. We hope to determine the effect of smoking e-cigarettes on the urine proteome of rats.

**Figure 1.**
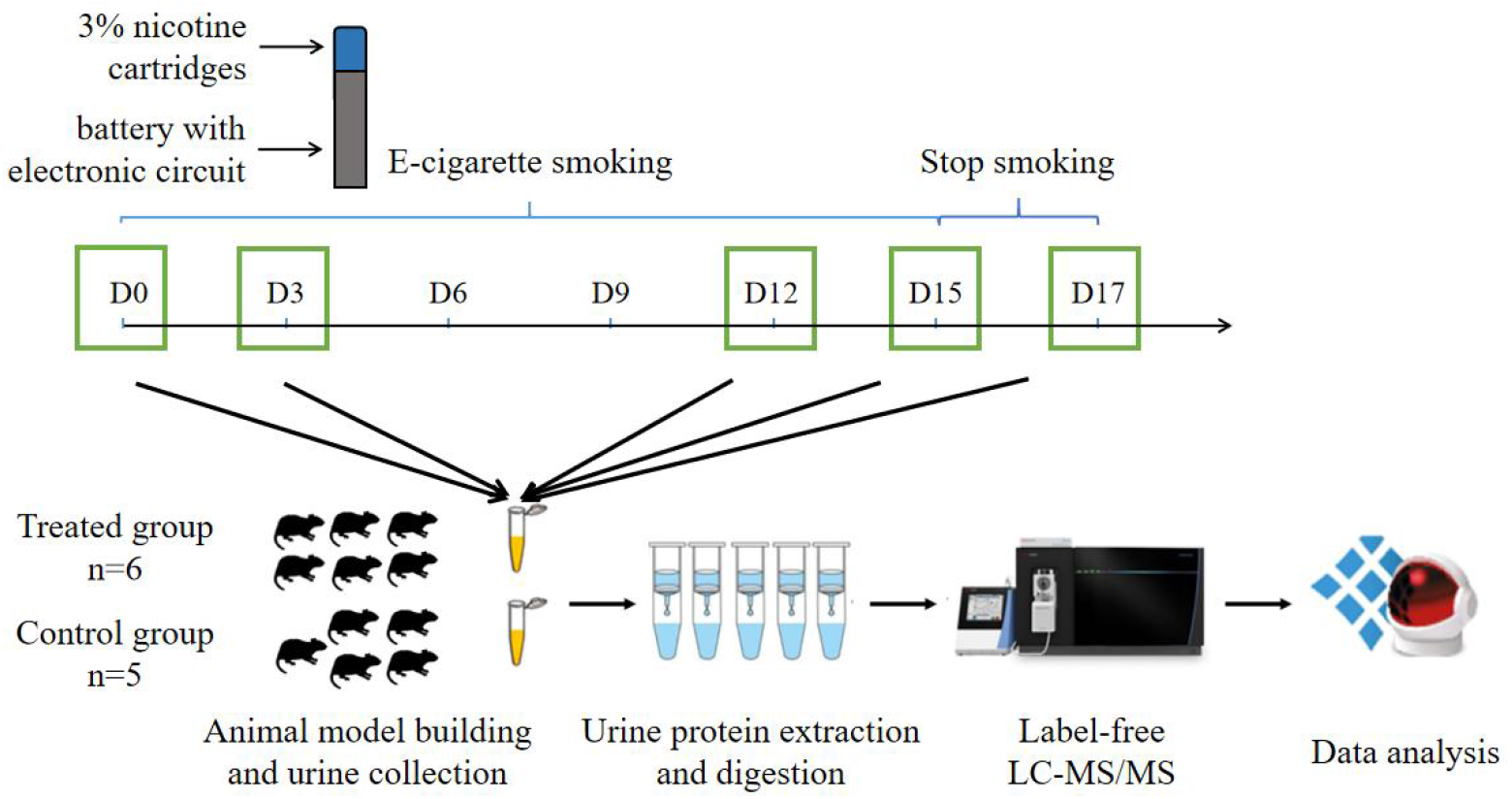
Workflow for urine proteomic analysis in rat e-cigarette models. In the experimental group, urine samples were collected before and on days 3, 6, 9, and 12 after smoking e-cigarettes and on days 1 and 3 after stopping smoking e-cigarettes. After urine samples were collected and processed in the experimental and control groups, the protein groups of the two groups were identified using liquid chromatography coupled with tandem mass spectrometry (LC–MS/MS) to quantitatively analyze the damage caused to the rat body at different stages of smoking e-cigarettes.

## 2. Materials and Methods

### 2.1 Rat model establishment

Eleven SPF 8-week-old healthy male Wistar rats (180-200 g) were purchased from Beijing Vital River Laboratory Animal Technology Co., Ltd., with the animal licence number SYXK (Jing) 2021-0011. All rats were maintained in a standard environment (room temperature (22±1)°C, humidity 65%–70%). All rats were kept in a new environment for three days before starting the experiment, and all experimental procedures followed the review and approval of the Ethics Committee of the College of Life Sciences, Beijing Normal University.

The animal model of e-cigarettes was established by randomly dividing 11 rats into an experimental group and a control group. Five of the control rats were maintained in a standard environment for 17 days. Six rats in the experimental group smoked e-cigarettes once a day during the same time period, and one-third of the 3% nicotine smoke bullets were made into smoke (approximately 16 mg of nicotine) and each time evenly injected into two cages [36 cm (length) × 20 cm (width) × 28 cm (height)]. And under the condition of ensuring adequate oxygen content, three rats in the experimental group were placed in each cage for 1 h and continued to smoke for 14 days, and they were returned to their cages after the end of smoking. Rats were observed for behavioural changes during the experiment, and body weights were recorded every 5 days.

### 2.2 Urine collection

After all rats were kept in a new environment for three days, they were uniformly placed in metabolic cages to collect urine samples for 12 h. Urine samples were collected for 12 h in metabolic cages from all rats on days 3, 6, 9, and 12 of e-cigarette smoking and on days 1 (as day 15) and 3 (as day 17) of cessation of e-cigarette smoking. Rats were fasted and water-deprived during urine collection, and all collected urine samples were stored in a −80 °C freezer.

### 2.3 Treatment of the urine samples

We wanted to observe the sensitivity of the urine proteome and observe whether the changes in the rat body could be reflected in the urine proteome after smoking e-cigarettes for a short time. Therefore, nonsmoking day 0, smoking days 3 and 12, and smoking cessation days 1 and 3 were selected as the samples for this focused analysis.

Urine protein extraction and quantification: Rat urine samples collected at five time points were centrifuged at 12,000×g for 40 min at 4 °C, and the supernatants were transferred to new Eppendorf (EP) tubes. Three volumes of precooled absolute ethanol were added, homogeneously mixed and precipitated overnight at −20 °C. The following day, the mixture was centrifuged at 12,000×g for 30 min at 4 °C, and the supernatant was discarded. The protein pellet was resuspended in lysis solution (containing 8 mol/L urea, 2 mol/L thiourea, 25 mmol/L dithiothreitol, 50 mmol/L Tris). The samples were centrifuged at 12,000×g for 30 min at 4 °C, and the supernatant was placed in a new EP tube. The protein concentration was measured by the Bradford assay.

Urinary protease cleavage: A 100 μg urine protein sample was added to the filter membrane (Pall, Port Washington, NY, USA) of a 10 kDa ultrafiltration tube and placed in an EP tube, and 25 mmol/L NH_4_HCO_3_ solution was added to make a total volume of 200 μL. Then, 20 mM dithiothreitol solution (dithiothreitol, DTT, Sigma) was added, and after vortex-mixing, the metal bath was heated at 97 °C for 5 min and cooled to room temperature. Iodoacetamide (Iodoacetamide, IAA, Sigma) was added at 50 mM, mixed well and allowed to react for 40 min at room temperature in the dark. Then, membrane washing was performed: ① 200 μL of UA solution (8 mol/L urea, 0.1 mol/L Tris-HCl, pH 8.5) was added and centrifuged twice at 14,000×g for 5 min at 18 °C; ② Loading: freshly treated samples were added and centrifuged at 14,000×g for 40 min at 18 °C; ③ 200 μL of UA solution was added and centrifuged at 14,000×g for 40 min at 18 °C, repeated twice; ④ 25 mmol/L NH_4_HCO_3_ solution was added and centrifuged at 14,000×g for 40 min at 18 °C, repeated twice; ⑤ trypsin (Trypsin Gold, Promega, Trypchburg, WI, USA) was added at a ratio of 1:50 of trypsin: protein for digestion and kept in a water bath overnight at 37 °C. The following day, peptides were collected by centrifugation at 13,000×g for 30 min at 4 °C, desalted through an HLB column (Waters, Milford, MA), dried using a vacuum dryer, and stored at −80 °C.

### 2.4 LC−MS/MS analysis

The digested samples were reconstituted with 0.1% formic acid, and peptides were quantified using a BCA kit, diluting the peptide concentration to 0.5 μg/μL. Mixed peptide samples were prepared from 4 μL of each sample and separated using a high pH reversed-phase peptide separation kit (Thermo Fisher Scientific) according to the instructions. Ten effluents (fractions) were collected by centrifugation, dried using a vacuum dryer and reconstituted with 0.1% formic acid. iRT reagent (Biognosys, Switzerland) was added at a volume ratio of sample:iRT of 10:1 to calibrate the retention times of extracted peptide peaks. For analysis, 1 μg of each peptide from an individual sample was loaded onto a trap column and separated on a reverse-phase C18 column (50 μm×150 mm, 2 μm) using the EASY-nLC1200 HPLC system (Thermo Fisher Scientific, Waltham, MA). The elution for the analytical column lasted 120 min with a gradient of 5%–28% buffer B (0.1% formic acid in 80% acetonitrile; flow rate 0.3 μL/min). Peptides were analyzed with an Orbitrap Fusion Lumos Tribrid Mass Spectrometer (Thermo Fisher Scientific, MA).

To generate the spectrum library, 10 isolated fractions were subjected to mass spectrometry in data-dependent acquisition (DDA) mode. Mass spectrometry data were collected in high sensitivity mode. A complete mass spectrometric scan was obtained in the 350-1500 m/z range with a resolution set at 60,000. Individual samples were analyzed using Data Independent Acquisition (DIA) mode. DIA acquisition was performed using a DIA method with 36 windows. After every 10 samples, a single DIA analysis of the pooled peptides was performed as a quality control.

### 2.5 Database searching and label-free quantitation

Raw data collected from liquid chromatography−mass spectrometry were imported into Proteome Discoverer (version 2.1, Thermo Scientific) and the Swiss-Prot rat database (published in May 2019, containing 8086 sequences) for alignment, and iRT sequences were added to the rat database. Then, the search results were imported into Spectronaut Pulsar (Biognosys AG, Switzerland) for processing and analysis. Peptide abundances were calculated by summing the peak areas of the respective fragment ions in MS_2_. Protein intensities were summed from their respective peptide abundances to calculate protein abundances.

### 2.6 Statistical analysis

Two technical replicates were performed for each sample, and the average was used for statistical analysis. In this experiment, the experimental group samples at different time periods were compared before and after, and the control group was set up to rule out differences in growth and development. Identified proteins were compared to screen for differential proteins. Differential protein screening conditions were as follows: fold change (FC) ≥ 1.5 or ≤ 0.67 between groups and *P* value < 0.05 by two-tailed unpaired t-test analysis. The Wukong platform was used for the selected differential proteins (Fig. https://www.omicsolution.org/wkomic/main/); the Uniprot website (Fig. https://www.uniprot.org/) and the DAVID database (Fig. https://david.ncifcrf.gov/) were used to perform functional enrichment analysis. In the PubMed database (Fig. https://pubmed.ncbi.nlm.nih.gov), the reported literature was searched to perform functional analysis of differential proteins.

## 3. Results

### 3.1 Characterization of e-cigarette smoking rats

In this experiment, the rats were observed behaviourally during the modeling process. Among them, rats in the control group had normal activity, a normal diet and drinking water. Water intake was significantly increased in the treated group compared with the control group. At the same time, the body weight of the rats was recorded every 5 days in this experiment (Fig. 2), and a significant increase in individualized differences in the body weight of the rats in the experimental group was observed.

**Figure 2.**
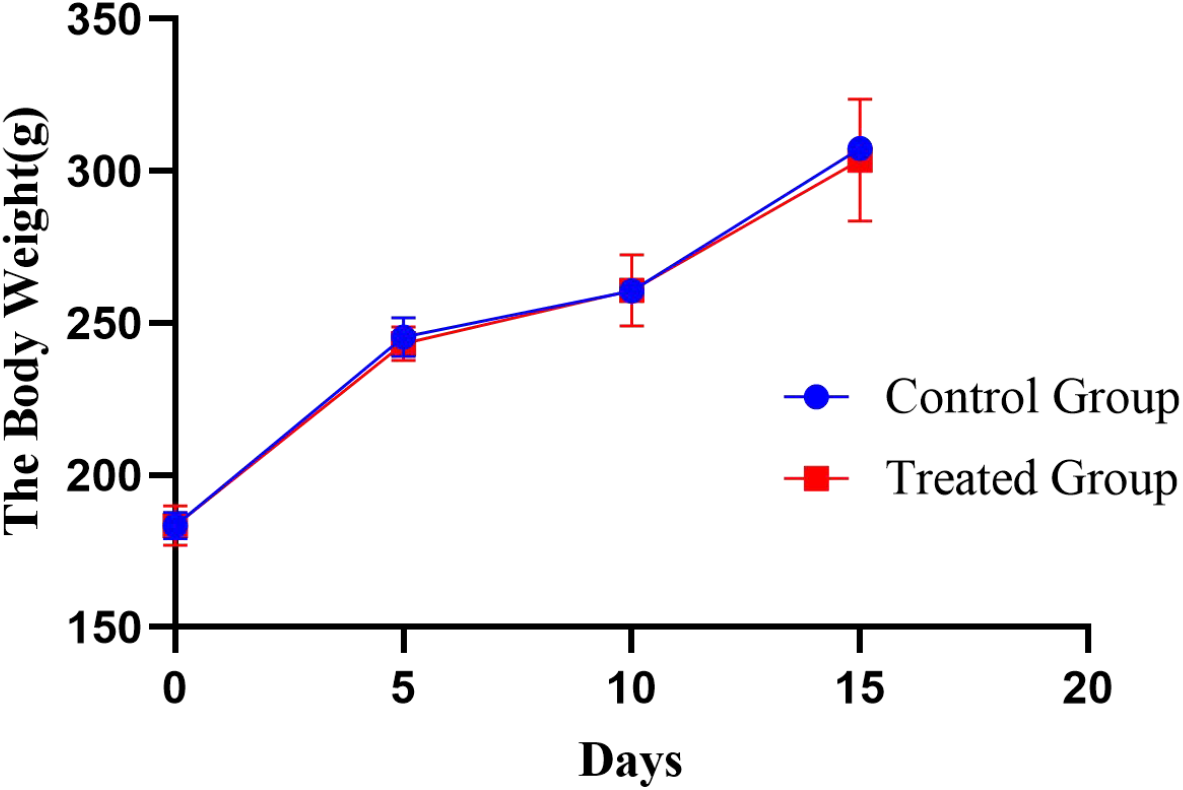
Body weight changes in the rat e-cigarette model. The obtained results are shown as the means±SDs for the control group (n=5) and the treated group (n=6).

### 3.2 Urinary proteome changes

#### 3.2.1 Urine protein identification

Fifty-five urine protein samples were analyzed by LC−MS/MS tandem mass spectrometry after the rat e-cigarette model was established. In total, 1093 proteins were identified (≥ 2 specific peptides and FDR < 1% at the protein level).

#### 3.2.2 Urine proteome changes

With the aim of investigating whether the changes were consistent in the six treated rats, we performed urine proteomics analysis of each rat individually for its own control and compared them at D0 at different time points. Screening differential protein conditions were FC ≥ 1.5 or ≤ 0.67, two-tailed unpaired t-test *P*<0.05. The differential protein screening results are shown in Table 1.

**Table 1.** Changes in differential protein expression during e-cigarette smoking in individual rats

| Days | Differential protein expression | Control group |  |  |  |  | Treated group |  |  |  |  |  |
| --- | --- | --- | --- | --- | --- | --- | --- | --- | --- | --- | --- | --- |
|  |  | Rat | Rat | Rat | Rat | Rat | Rat | Rat | Rat | Rat | Rat | Rat |
|  |  | 1 | 2 | 3 | 4 | 5 | 1 | 2 | 3 | 4 | 5 | 6 |
| Day3/Day0 | Total | 215 | 208 | 208 | 162 | 96 | 181 | 203 | 177 | 331 | 144 | 490 |
|  | ↑ | 89 | 83 | 82 | 71 | 59 | 51 | 87 | 58 | 127 | 60 | 199 |
|  | ↓ | 126 | 125 | 126 | 91 | 37 | 130 | 116 | 119 | 204 | 84 | 291 |
| Day12/Day0 | Total | 401 | 257 | 214 | 275 | 217 | 260 | 265 | 188 | 336 | 173 | 533 |
|  | ↑ | 175 | 115 | 82 | 126 | 106 | 117 | 112 | 80 | 152 | 93 | 241 |
|  | ↓ | 226 | 142 | 132 | 149 | 111 | 143 | 153 | 108 | 184 | 80 | 292 |
| Day15/Day0 | Total | 288 | 255 | 337 | 289 | 505 | 350 | 341 | 261 | 402 | 246 | 512 |
|  | ↑ | 124 | 141 | 130 | 104 | 312 | 125 | 153 | 104 | 163 | 151 | 214 |
|  | ↓ | 173 | 114 | 207 | 185 | 193 | 225 | 188 | 157 | 239 | 95 | 298 |
| Day17/Day0 | Total | 395 | 264 | 289 | 320 | 191 | 266 | 283 | 199 | 290 | 475 | 518 |
|  | ↑ | 183 | 134 | 111 | 157 | 142 | 117 | 140 | 81 | 124 | 312 | 251 |
|  | ↓ | 212 | 130 | 178 | 163 | 49 | 149 | 143 | 188 | 166 | 163 | 267 |

To observe the consistency of the changes in the six treated rats more intuitively, we made Venn diagrams of the differential proteins screened by comparing the six rats in the experimental group before and after on D3, D12, D15, and D17 with those on D0 (Fig. 3).

**Figure 3.**
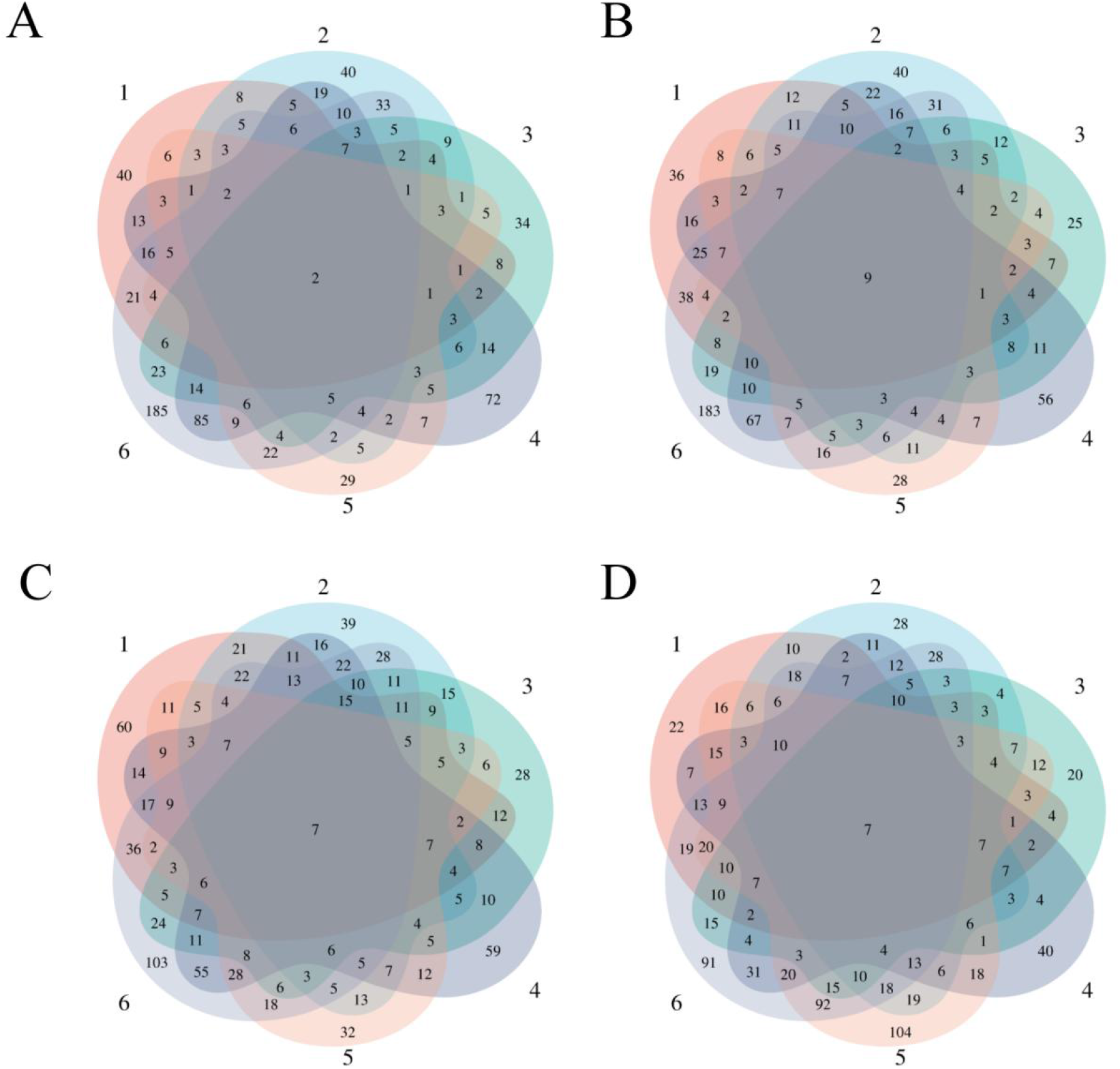
Differential proteins Venn diagram produced by 6 treated rats and their own control. (A) D3 vs. D0. (B) D12 vs. D0. (C) D15 vs. D0. (D) D17 vs. D0.

To verify the correlation between rat differential proteins at different time points, we made Venn diagrams. They show the common differential proteins identified in urine protein from five or more treated rats on D3, D12, D15, and D17 compared with D0 (Fig. 4). Among them, cadherin was screened by 5 or more rats at all four time points, and all showed a tendency of downregulation. Six differential proteins were screened by 5 or more rats at all three time points, showing more common differences. The trends of common differential proteins in the six rats of the treated group on different days are shown in Table 2.

**Table 2.**
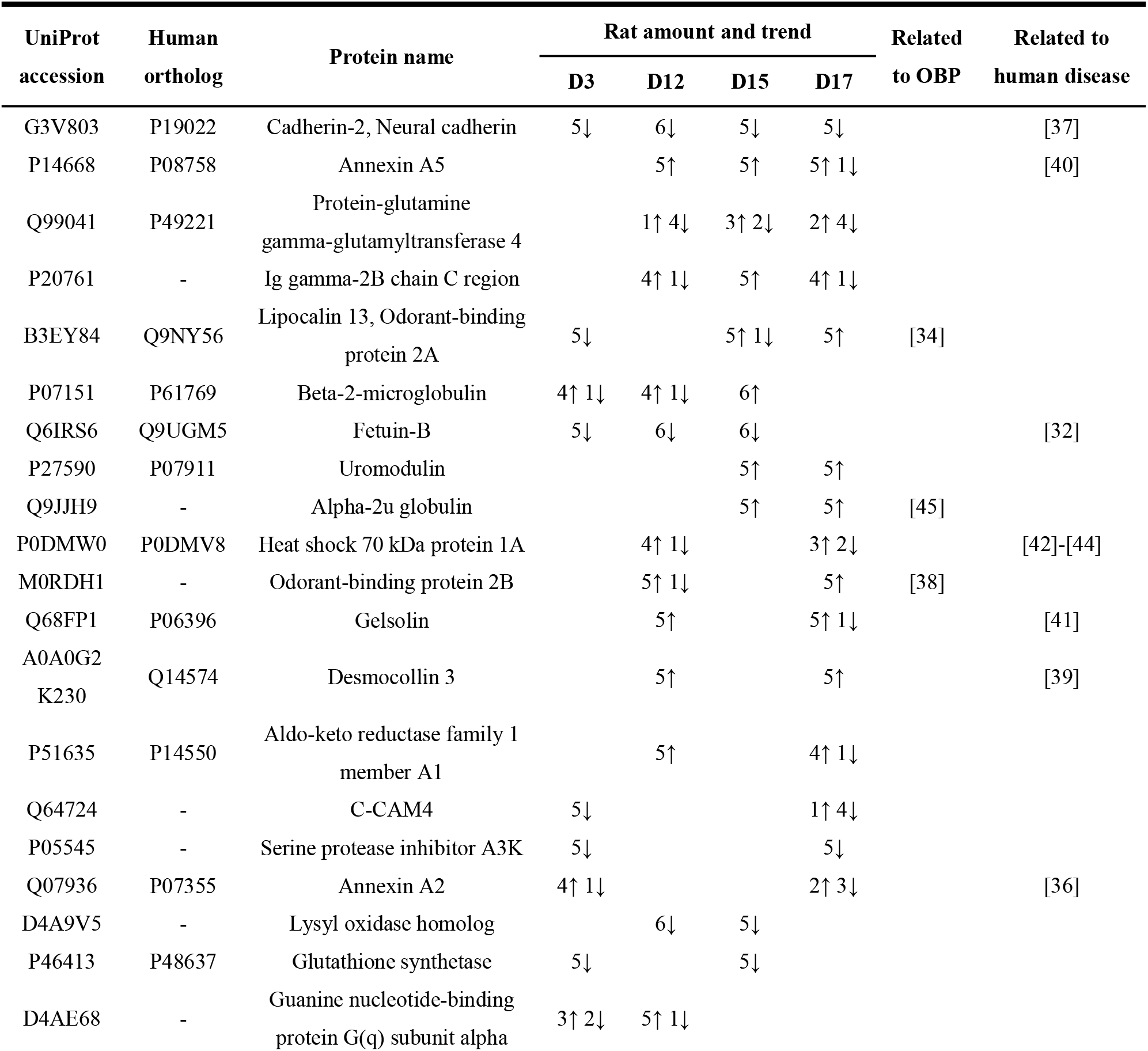
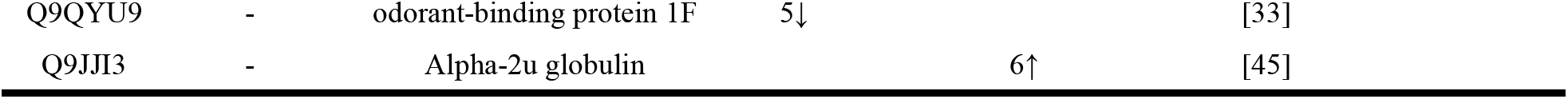
Trends of common differential proteins in 6 rats in the experimental group on different days

**Figure 4.**
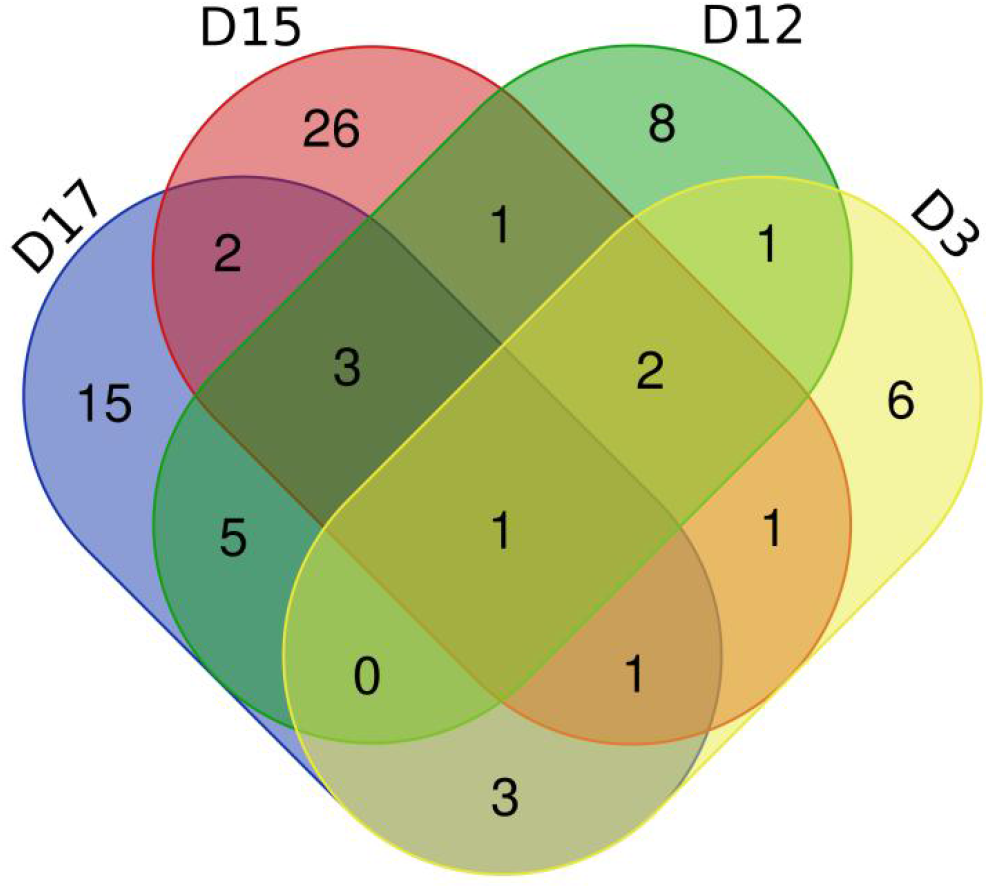
Venn diagram of common differential proteins in 6 rats in the experimental group on different days

We analyzed differential proteins produced by D3 after e-cigarette smoking and D0 self-controls in six treated rats. We found that two differential proteins were commonly identified in six treated rats, and 16 differential proteins were commonly identified in five experimental rats. After comparing the differential proteins produced by the control group before and after itself, the repeated proteins were screened out, resulting in differential proteins with specific commonalities in the treated group. The details are presented in Table S1.

We also analyzed the consensus differential proteins produced by six rats in the experimental group on D12 after e-cigarette smoking versus D0 self-control. Nine differential proteins were commonly identified in six treated rats, and 18 differential proteins were commonly identified in five experimental rats. After comparison with the differential proteins produced by a single rat in the control group before and after treatment, the repeated proteins were screened out, resulting in differential proteins with specific commonalities in the treated group. The details are presented in Table S2.

When D15 was compared with D0, 7 differential proteins were commonly identified in 6 treated rats, and 45 differential proteins were commonly identified in 5 experimental rats. After comparing the differential proteins produced by the control group before and after, the repeated proteins were screened out, resulting in differential proteins with specific commonalities in the experimental group. The details are presented in Table S3.

Finally, we analyzed six rats in the experimental group for differential proteins screened by self-comparison between D17 and D0, of which 7 differential proteins were commonly identified from 6 rats in the experimental group and 42 differential proteins were commonly identified from 5 rats in the experimental group. After comparing the differential proteins produced by the control group before and after, the repeated proteins were screened out, resulting in differential proteins with specific commonalities in the experimental group (see Table S4 for details). Among the results presented at different time points, we found some consistency in the effects caused by e-cigarette smoking in rats.

#### 3.2.3 Functional comparison analysis

To investigate the function of these differential proteins, we performed functional analysis of biological pathways using the DAVID database on the differential proteins coselected from five or more treated rats (see Table S5 for details). Thirty-two of these biological processes were enriched by differential proteins at both time points. We also performed signaling pathway analysis of differentially identified proteins coidentified in five or more treated rats (Table 3). Two of these signaling pathways were coenriched at both time points. These include Legionellosis and Ferroptosis. In addition, we also enriched many signaling pathways associated with respiratory diseases, such as the apelin signaling pathway^[28][29]^, folate biosynthesis pathway^[30]^, and arachidonic acid metabolism^[31]^. Of note, we also found two signaling pathways associated with chemical carcinogenesis, including chemical carcinogenesis-DNA adducts and chemical carcinogenesis-reactive oxygen species. This finding reinforces the sensitivity of the urine proteome.

**Table 3.**
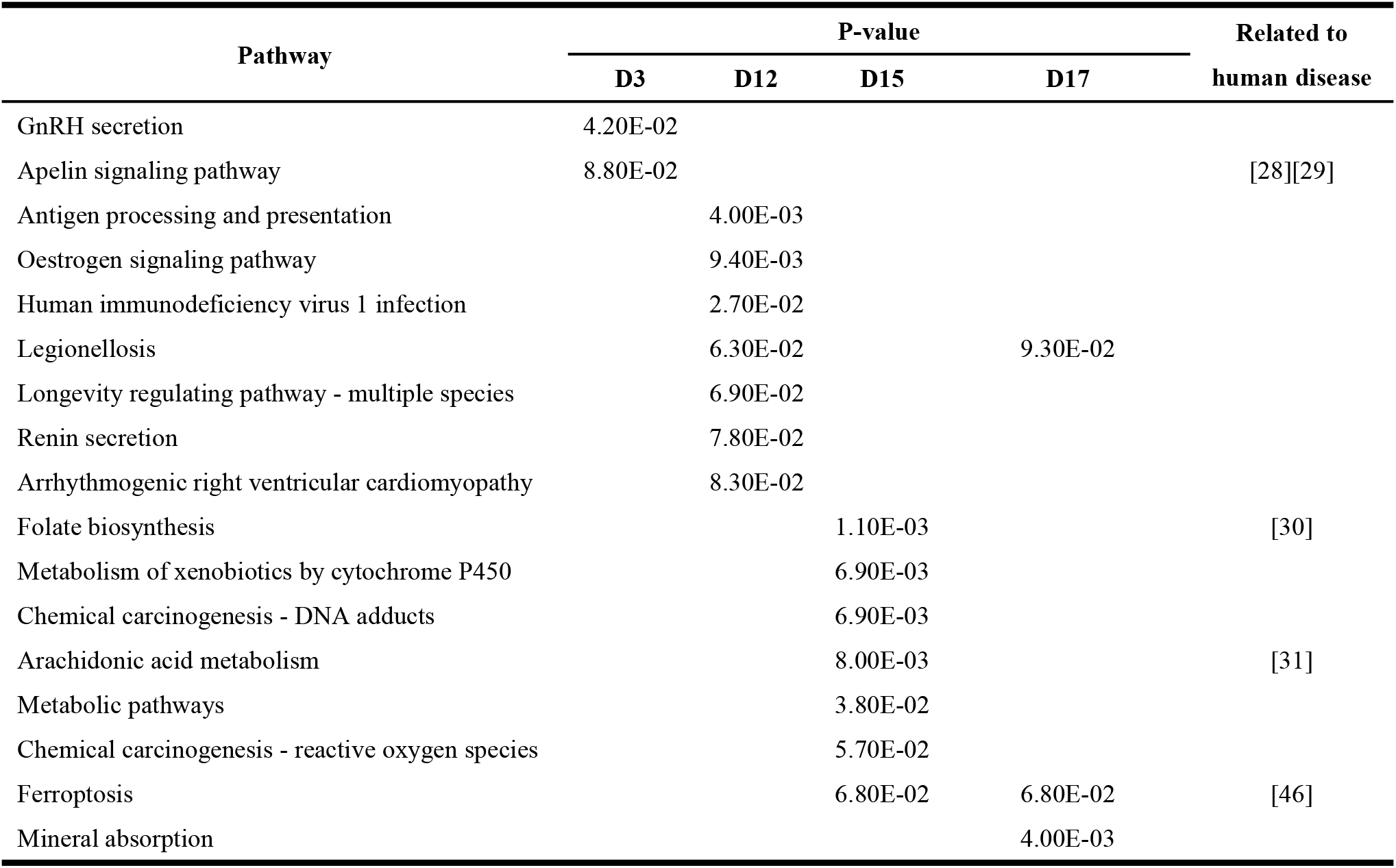
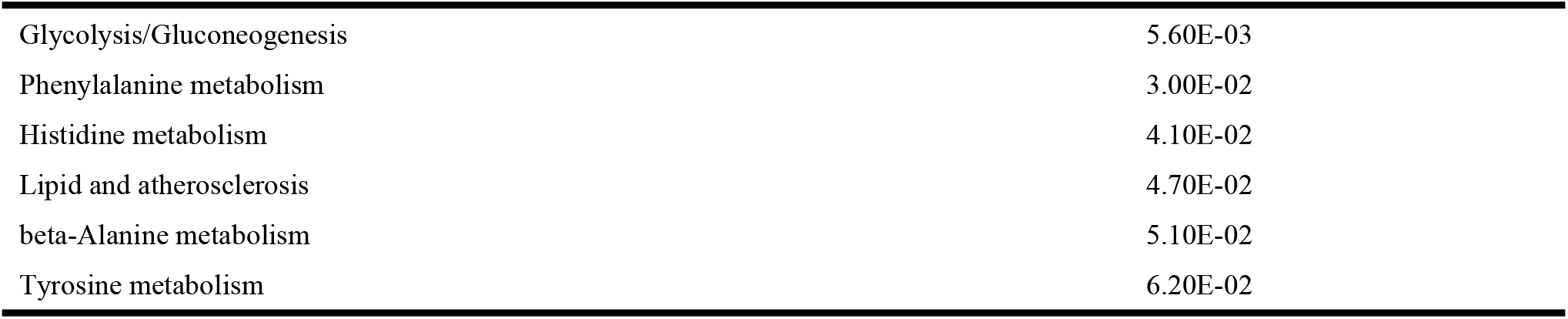
Signal pathways enriched in common differential proteins produced during e-cigarette smoking in five or more treated rats.

## 4. Discussion

In this study, we constructed a rat e-cigarette model and collected urine samples before, during, and after e-cigarette smoking in rats on days 0, 3, 12, 15, and 17 to explore e-cigarettes from the perspective of urine proteomics. To exclude the influence of individual differences, the experiment used a single rat before and after the control for analysis, while the control group was set up to rule out differences caused by rat growth and development. From the results presented by the Venn diagram (Fig. 3) of the differential proteins screened from 6 rats in the experimental group on days 3, 12, 15, and 17 compared with themselves on day 0, we found that most of the differential proteins were personalized, indicating that the effects caused by e-cigarettes on rats had strong individualized differences.

We analyzed six treated rats D3 compared with D0; among the resulting differential proteins, fetuin-B was identified in five rats, all of which showed a significant decreasing trend. There were also significant differences in urine protein on D12 and D15. It has been shown that fetuin-B is a biomarker of chronic obstructive pulmonary disease (COPD)^[32]^, reflecting the sensitivity of the urine proteome. Surprisingly, we also observed odorant-binding protein (OBP), including OBP1F and OBP2A, in urine protein with significant changes on D3, of which OBP1F is mainly expressed in the nasal glands of rats^[33]^, and OBP2A is also mainly transcribed in the nose of humans and rats^[34]^. This suggests that rat smoking odorants can actually leave traces in the urine proteome. Currently, the physiological role of OBPs is not fully understood^[35]^, and perhaps the urine proteome can play a role in exploring the specific mechanism of action of OBPs. In addition, OBPs appear in the urine proteome, which may provide an explanation for olfactory adaptation. Annexin A2 is widely used as a marker for a variety of tumors^[36]^. In addition, László ZI et al. showed that neurocadherin is one of the most important cell adhesion molecules during brain development and plays an important role in neuronal formation, neuronal proliferation, differentiation and migration, axonal guidance, synaptogenesis and synaptic maintenance^[37]^.

We also compared D12 with D0 of six rats in the experimental group. Among the differential proteins produced in all six treated rats, OBP2B also showed a more consistent upregulation. Unlike OBP2A, OBP2B is mainly expressed in reproductive organs and weakly expressed in organs of the respiratory system such as the nose and lung^[38]^. In addition, it has been shown that desmocollin 3 is an essential protein for cell adhesion and desmosome formation and may enhance angiogenesis with metastasis in nasopharyngeal carcinoma and is considered a biomarker for some cancers, such as non-small cell lung cancer^[39]^. It has also been shown that Annexin A5 may affect the occurrence and development of pathological phenomena, such as tumor diseases, pulmonary fibrosis and lung injury. It may also be used as a biomarker in the study of diseases, such as tumors and asthma, and may also promote the occurrence and development of laryngeal cancer and nasopharyngeal carcinoma^[40]^. We also screened an important cellular target of the nicotine metabolite cotinine, gelsolin, which may affect basic processes of tumor transformation and metastasis, such as migration and apoptosis, through gelatin^[41]^. Heat shock protein, on the other hand, is reported to be a major marker affected by cigarette smoke and is involved in signaling pathways for the cell cycle and cell death and inflammation^[42]-[44]^.

Comparing D15 with D0, the common differential proteins produced by six experimental rats contained α-2u globulin. Ponmanickam P et al. showed that α-2u globulin may act as a carrier of hydrophobic odorants in the preputial gland, which plays an important role in producing pheromone-communicating olfactory signals in rats. Therefore, α-2u globulin is likely to be involved in the transmission of olfactory signals in rats^[45]^.

Compared with D0, most of the common differential proteins produced by six rats in the experimental group on D17 were similar to those produced on other days, which all contain odorant-binding proteins and proteins associated with a variety of diseases (Table 2).

We performed signaling pathway analysis of differentially expressed proteins coidentified in five or more treated rats (Table 3) and focused on two signaling pathways coenriched by the two time points: Legionellosis and Ferroptosis. Pneumonia caused by Legionnaires’ disease may cause damage to the body similar to the mechanism of the effects of smoking e-cigarettes on the body. Whereas M. Yoshida et al. showed that smoking can induce Ferroptosis in epithelial cells, and this signaling pathway is involved in the pathogenesis of chronic obstructive pulmonary disease^[46]^. In addition, we also enriched many signaling pathways associated with respiratory diseases, such as the apelin signaling pathway^[28][29]^, folate biosynthesis pathway^[30]^, and arachidonic acid metabolism^[31]^. Apelin is an endogenous ligand for the G protein-coupled receptor APJ^[28]^, and the apelin/APJ pathway is closely related to the development of respiratory diseases. Targeting the apelin/APJ system may be an effective therapeutic approach for respiratory diseases^[29]^. A study by Staniszavska-Sachadyn A et al. showed that serum folate concentrations were higher in smokers than in healthy controls, and it was postulated that folate synthesis is associated with an increased risk of lung cancer^[30]^. Because the 23 enriched signaling pathways include multiple signaling pathways directly related to the immune system, cardiomyopathy, and atherosclerosis, we speculated that smoking e-cigarettes may affect the immune system and cardiovascular system of rats. For the two signaling pathways enriched to be associated with chemical carcinogenesis, it may be possible to verify previous findings that e-cigarette smoke contains carcinogenic chemicals^[47]^.

## 5. Conclusion

There were strong individual differences in the differential proteins produced by rats after smoking e-cigarettes under the same conditions. Fetuin-B, a biomarker of COPD, and annexin A2, which is recognized as a multiple tumor marker, were co-identified in five out of six treated rats’ self-control on D3. Odorant-binding proteins expressed in the olfactory epithelium were also identified in the urine proteome at multiple time points and were significantly upregulated, which may help explain olfactory adaptation. How do odorant binding proteins expressed in the olfactory epithelium end up in the urine after smoking e-cigarettes remain to be elucidated. The evidences that smoking e-cigarettes affects the immune system, cardiovascular system, respiratory system, etc. were found in both the differential proteins and the enriched signaling pathways, providing clues for further exploration of the mechanism of e-cigarettes on the human body.

## CRediT authorship contribution statement

**Yuqing Liu:** Conceptualization, Investigation, Writing-original draft, Data curation, Visualization. **Ziyun Shen:** Investigation. **Chenyang Zhao:** Data curation. **Youhe Gao:** Conceptualization, Writing-review & editing, Supervision, Funding acquisition.

## Acknowledgements

This work was supported by the National Key Research and Development Program of China (2018YFC0910202); the Fundamental Research Funds for the Central Universities (2020KJZX002); the Beijing Natural Science Foundation (7172076).

---

**Table S1.** Differential proteins identified in the D3 test group before and after self-control in 6 rats

| UniProt<br>accession | Human<br>ortholog | Protein name | Fold change |  |  |  |  |  |
| --- | --- | --- | --- | --- | --- | --- | --- | --- |
|  |  |  | Rat<br>1 | Rat<br>2 | Rat<br>3 | Rat<br>4 | Rat<br>5 | Rat<br>6 |
| P08721 | P10451 | Osteopontin | 3.29 | 3.60 | 2.93 | 0.65 | 2.07 | 0.55 |
| A0A0G2JYF9 | O15232 | Matrilin 3 | 4.00 | 2.11 | 2.24 | 2.93 | 2.51 | 0.44 |
| P07151 | P61769 | Beta-2-microglobulin | 1.91 | 1.73 | 2.23 | 0.62 | 2.35 | - |
| Q6IRS6 | Q9UGM5 | Fetuin-B | 0.47 | 0.40 | 0.47 | 0.32 | 0.31 | - |
| Q9QYU9 | - | Odorant-binding protein 1F | 0.53 | 0.46 | 0.41 | 0.65 | - | 0.26 |
| Q64724 | - | C-CAM4 | 0.59 | 0.67 | 0.53 | 0.40 | - | 0.22 |
| B3EY84 | Q9NY56 | Lipocalin 13, odorant-binding protein 2A | 0.59 | 0.59 | 0.39 | 0.63 | - | 0.22 |
| P22006 | - | Seminal vesicle secretory protein 2 | 0.37 | 3.97 | 9.74 | 0.12 | - | 0.09 |
| F1LP05 | - | ATP synthase subunit alpha | 0.38 | 0.45 | 0.31 | - | 2.56 | 5.19 |
| Q07936 | P07355 | Annexin A2 | 0.60 | 4.19 | - | 1.59 | 1.82 | 2.39 |
| P05545 | - | Serine protease inhibitor A3K | 0.62 | 0.60 | - | 0.64 | 0.56 | 0.46 |
| D4AE68 | - | Guanine nucleotide-binding protein G(q) subunit alpha | - | 0.57 | 0.28 | 2.32 | 2.84 | 4.03 |
| G3V803 | P19022 | Cadherin-2, Neural cadherin | - | 0.65 | 0.56 | 0.59 | 0.57 | 0.32 |
| P46413 | P48637 | Glutathione synthetase | - | 0.42 | 0.50 | 0.42 | 0.52 | 0.45 |
| Q5I0D7 | P12955 | Xaa-Pro dipeptidase | - | 0.39 | 0.59 | 0.44 | 0.65 | 0.45 |

**Table S2.** Differential proteins identified in the D12 test group before and after self-control in 6 rat

| UniProt<br>accession | Human<br>ortholog | Protein name | Fold change |  |  |  |  |  |
| --- | --- | --- | --- | --- | --- | --- | --- | --- |
|  |  |  | Rat<br>1 | Rat<br>2 | Rat<br>3 | Rat<br>4 | Rat<br>5 | Rat<br>6 |
| D4AE68 | - | Guanine nucleotide-binding protein G(q) subunit alpha | 2.89 | 1.80 | 0.50 | 3.07 | 1.57 | 4.83 |
| Q6IRS6 | Q9UGM5 | Fetuin-B | 0.42 | 0.57 | 0.62 | 0.53 | 0.53 | 0.33 |
| M0RDH1 | - | Odorant-binding protein 2B | 2.49 | 2.20 | 2.14 | 2.90 | 2.17 | 0.46 |
| G3V803 | P19022 | Cadherin-2, Neural cadherin | 0.47 | 0.54 | 0.42 | 0.57 | 0.58 | 0.22 |
| D4A9V5 | - | Lysyl oxidase homolog | 0.54 | 0.40 | 0.67 | 0.47 | 0.41 | 0.20 |
| P07151 | P61769 | Beta-2-microglobulin | 2.75 | 2.07 | 2.84 | 0.57 | 2.08 | 2.19 |
| A0A0G2K230 | Q14574 | Desmocollin 3 | 1.96 | 2.73 | 4.04 | 2.41 | 3.02 | - |
| P51635 | P14550 | Aldo-keto reductase family 1 member A1 | 2.29 | 1.62 | 1.96 | 2.87 | 3.40 | - |
| P14668 | P08758 | Annexin A5 | 1.56 | 1.84 | 2.92 | 2.96 | 5.95 | - |
| G3V8V1 | - | Granulin, isoform CRA_c | 0.54 | 0.58 | 0.52 | - | 0.22 | 0.32 |
| A0A0G2K6Z6 | - | Tartrate-resistant acid phosphatase type 5 | 0.38 | 0.39 | 0.49 | - | 0.29 | 0.27 |
| P20761 | - | Ig gamma-2B chain C region | 0.33 | 1.79 | 1.78 | - | 1.69 | 2.55 |
| Q68FP1 | P06396 | Gelsolin | 2.20 | 1.82 | 2.10 | - | 2.51 | 2.38 |
| F1LPR6 | - | Immunoglobulin heavy constant epsilon | 2.67 | 2.85 | - | 2.02 | 2.94 | 3.33 |
| F1M5L5 | - | Ig-like domain-containing protein | 3.89 | 1.75 | - | 2.78 | 1.97 | 0.53 |
| P12020 | P54107 | Cysteine-rich secretory protein 1 | 2.38 | 2.41 | - | 0.13 | 4.77 | 4.95 |
| F1M8K0 | - | Dystroglycan 1 | 0.50 | 0.47 | - | 0.23 | 0.62 | 0.47 |
| Q6VPP3 | - | Chloride channel accessory 4 | 0.62 | 0.30 | - | 0.39 | 1.61 | 0.65 |
| G3V8T4 | - | DNA damage-binding protein 1 | 2.51 | 1.69 | - | 4.95 | 1.99 | 2.72 |
| Q99041 | P49221 | Protein-glutamine gamma-glutamyltransferase 4 | - | 0.05 | 0.04 | 0.06 | 0.24 | 16.03 |
| P0DMW0 | P0DMV8 | Heat shock 70 kDa protein 1A | - | 2.75 | 2.41 | 0.33 | 5.65 | 3.30 |

**Table S3.**
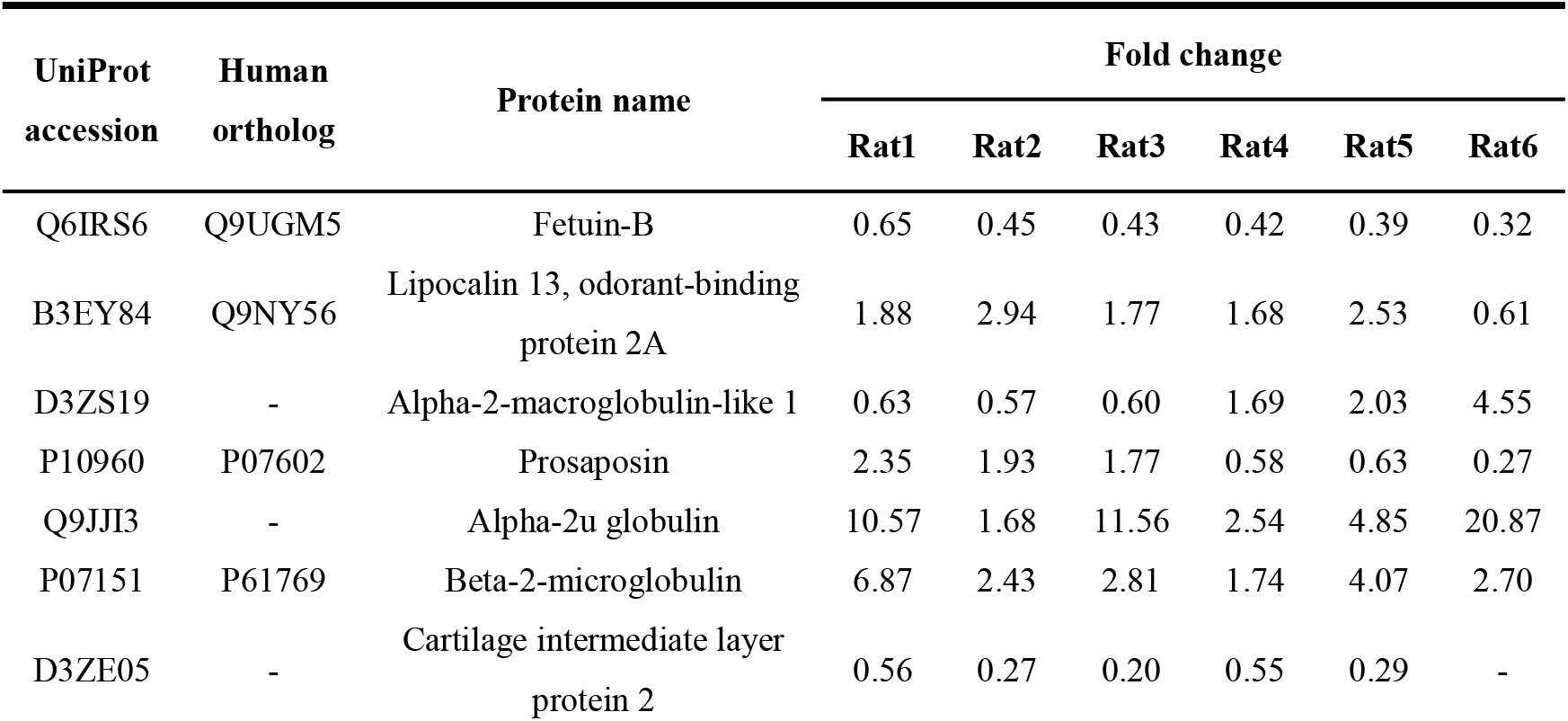
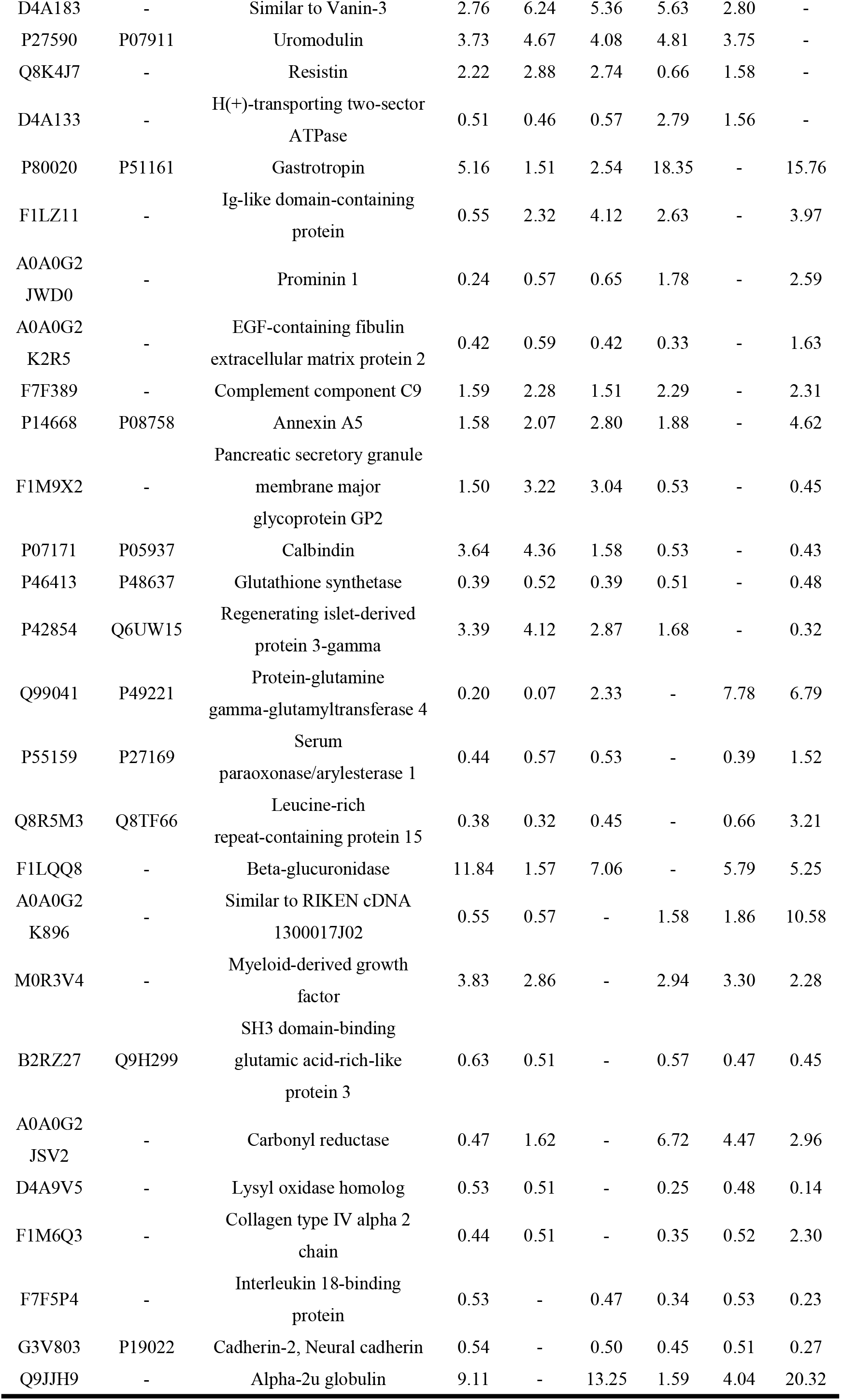
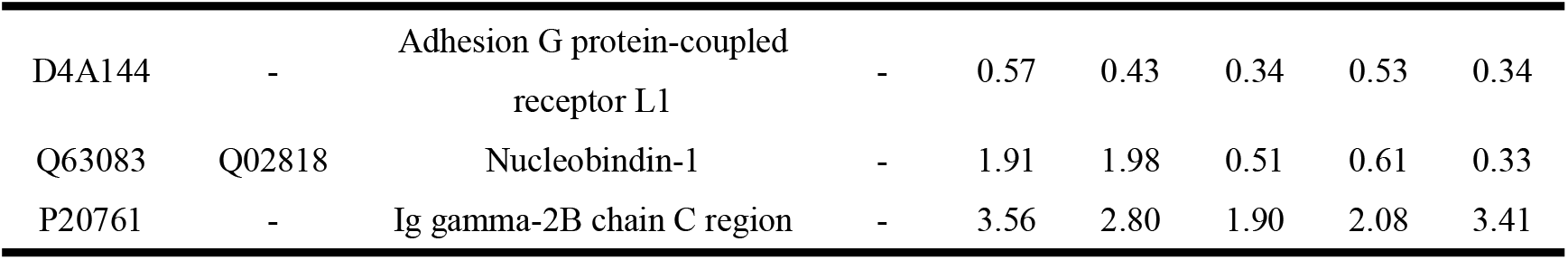
Differential proteins identified in the D15 test group before and after self-control in 6 rats

**Table S4.**
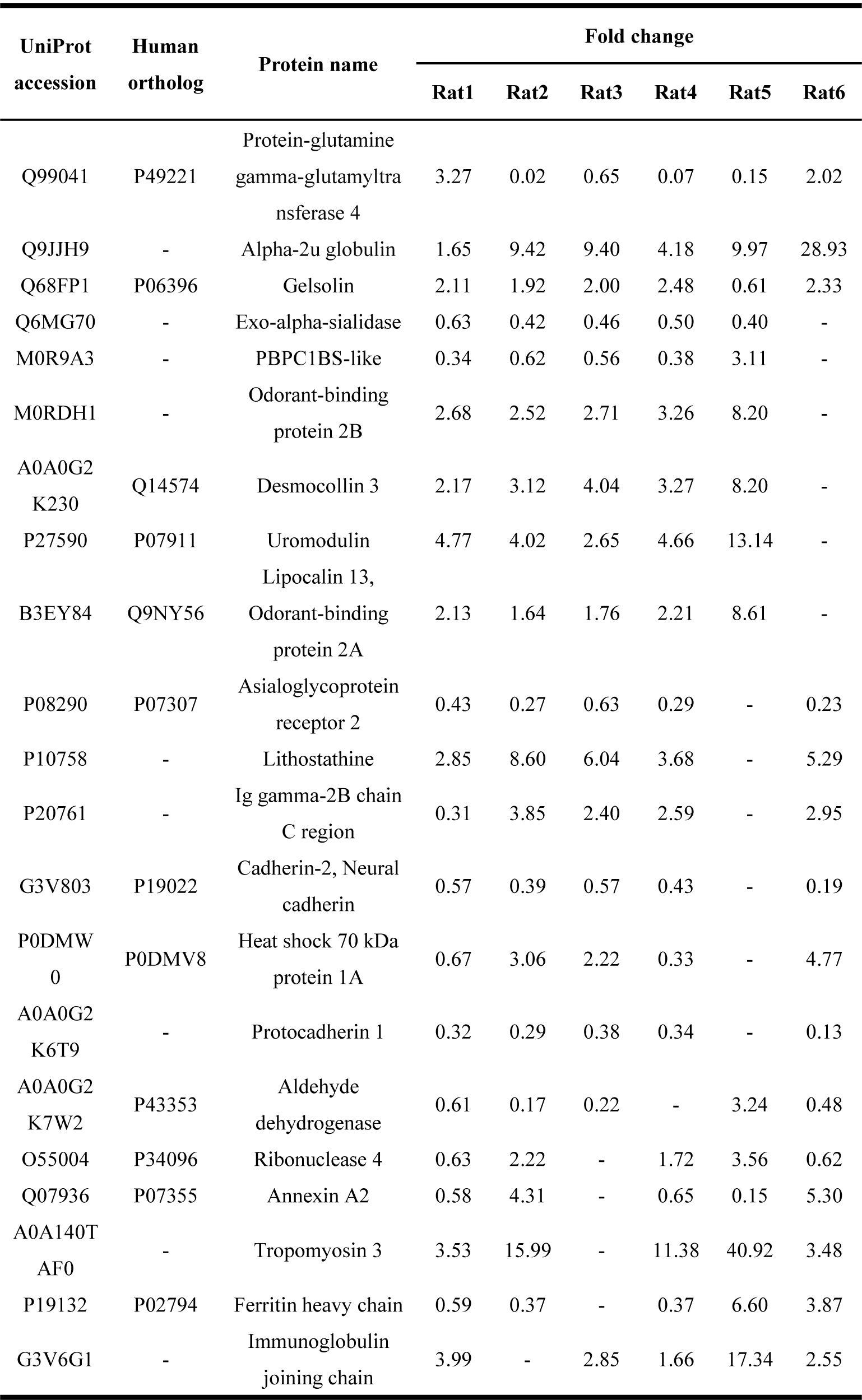
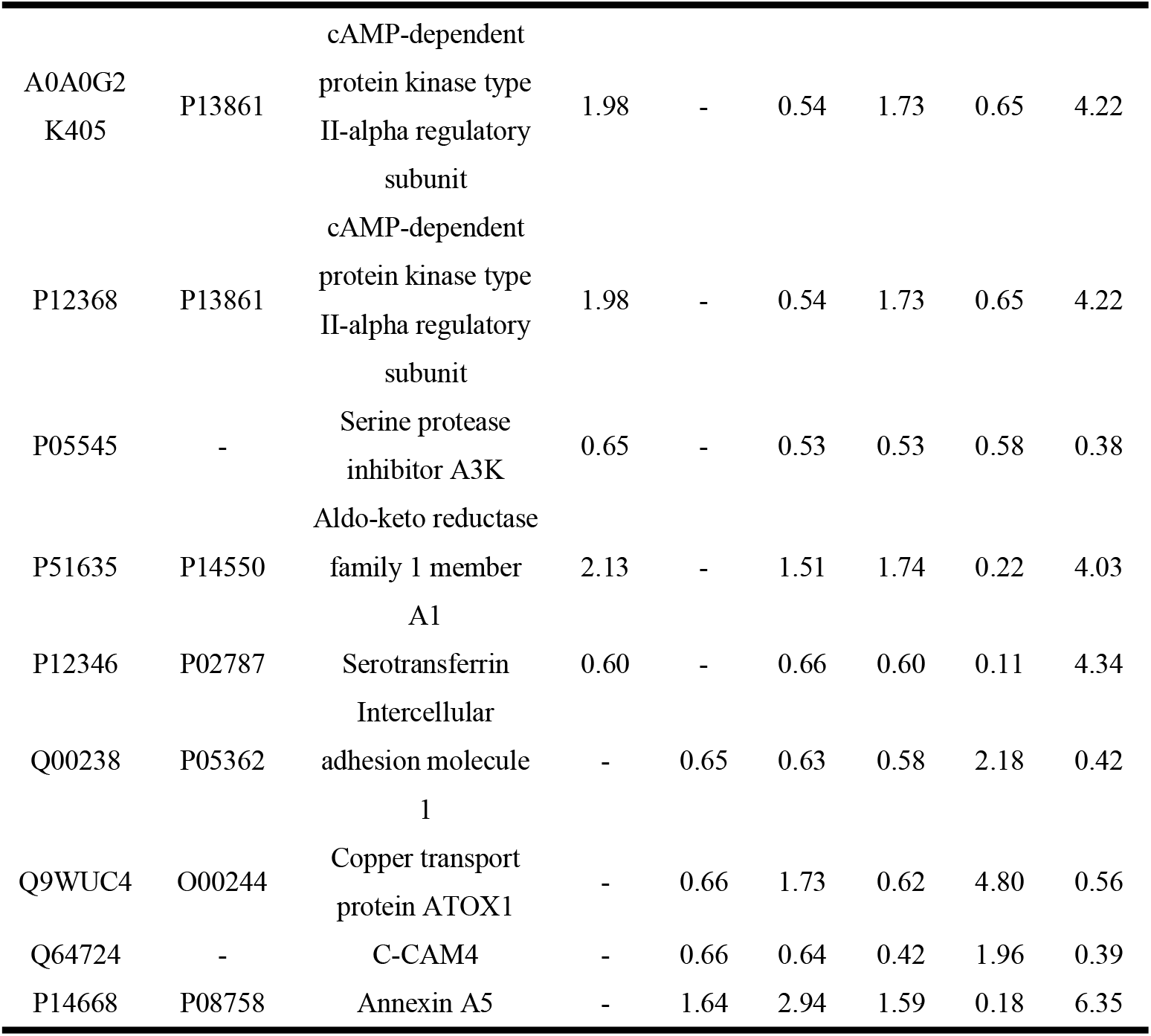
Differential proteins identified in the D17 test group before and after self-control in 6 rats

**Table S5.**
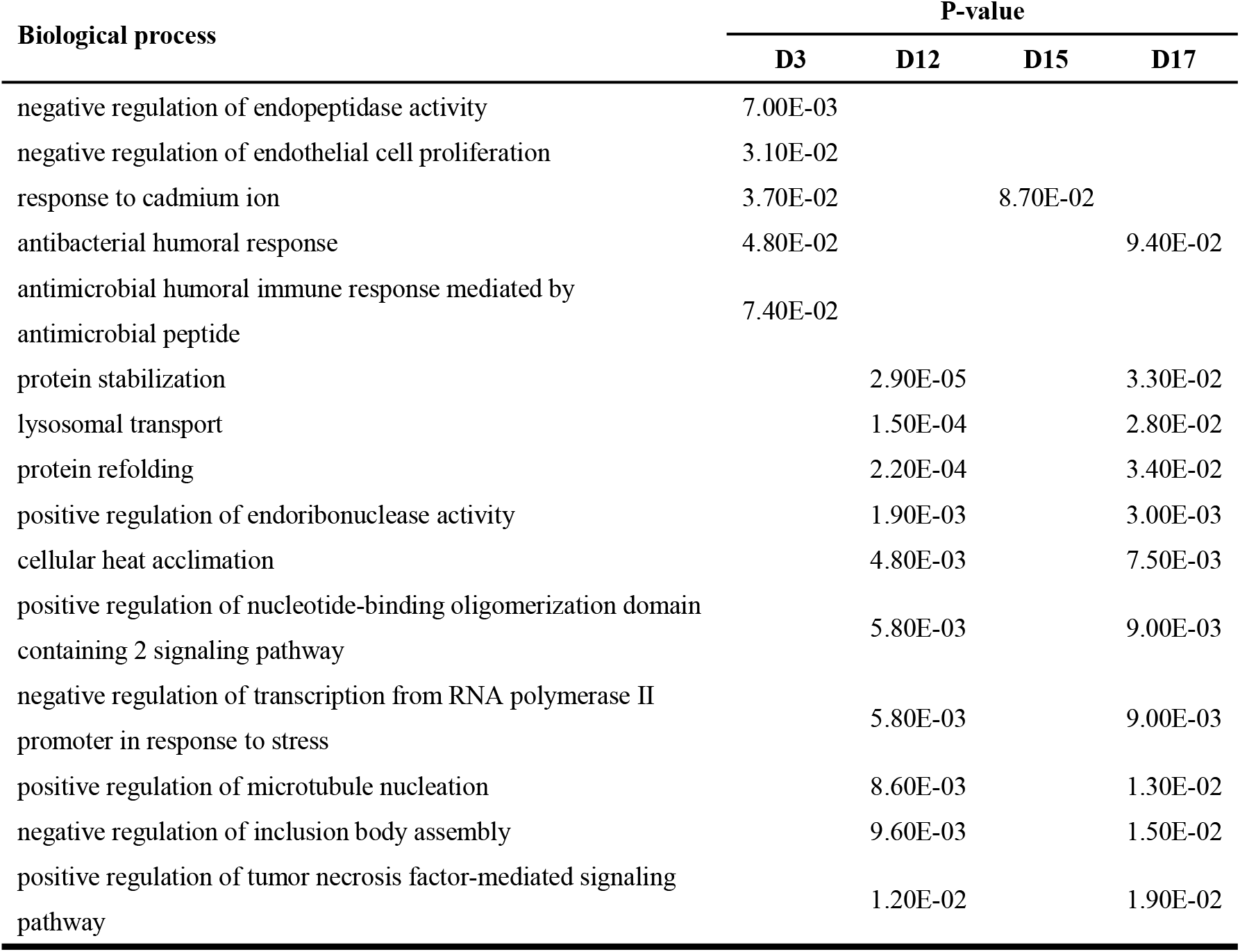
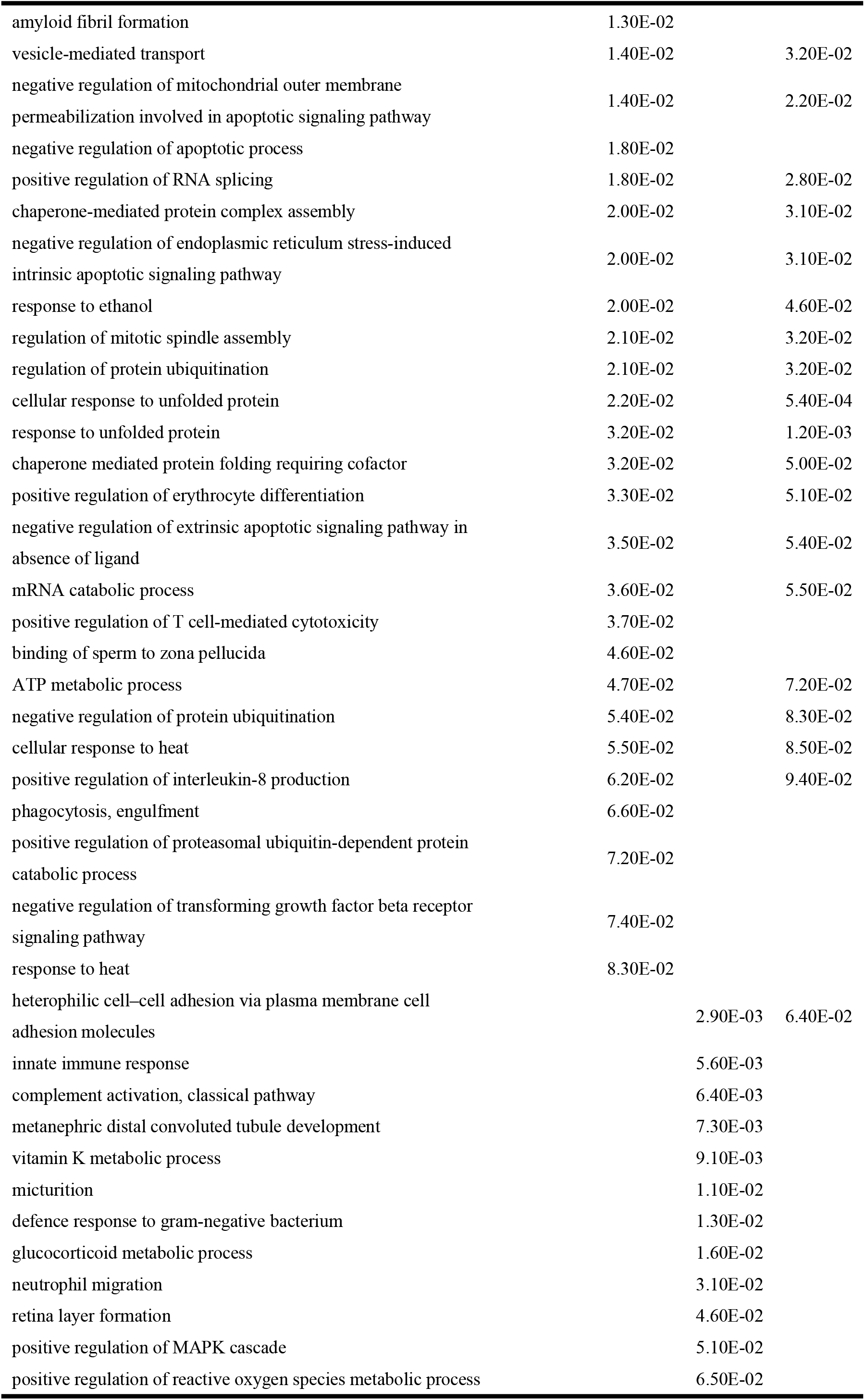
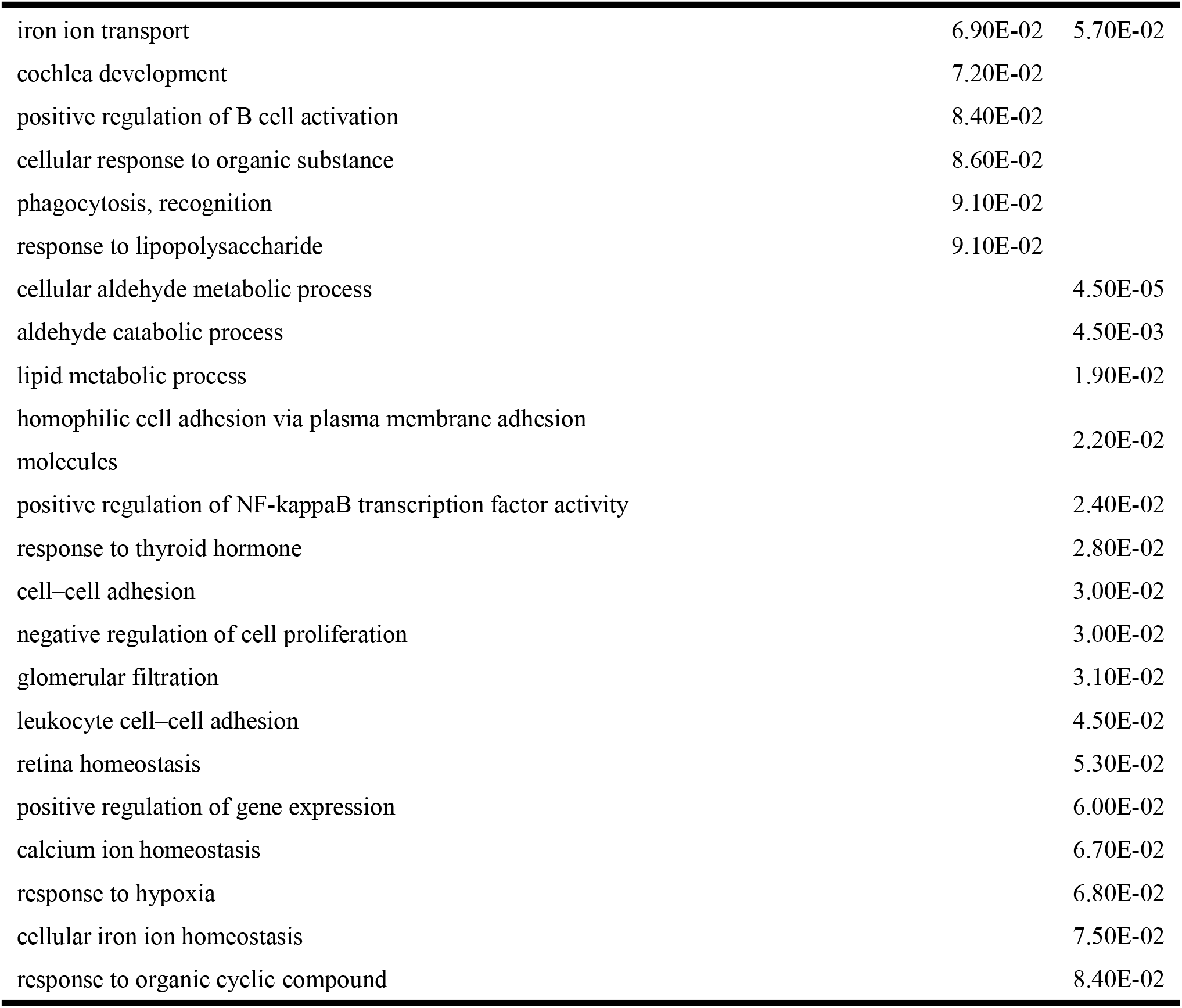
Biological processes enriched by the differential proteins identified jointly before and after comparison in five or more treated rats

## References

[1] http://www.xinhuanet.com/politics/2019-11/06/c_1125199903.htm

[2] Bold KW, Kong G, et al. Reasons for trying E-cigarettes and risk of continued use. Pediatrics. 2016, 138(3):e20160895.

[3] Pipe AL, Mir H. E-Cigarettes Reexamined: Product Toxicity. Can J Cardiol. 2022, 38(9):1395–1405.

[4] Cheng KA, Nichols H, McAdams HP, et al. Washington L. Imaging of Smoking and Vaping Related Diffuse Lung Injury. Radiol Clin North Am. 2022, 60(6):941–950.

[5] Fetterman JL, Keith RJ, Palmisano JN, et al. Alterations in vascular function associated with the use of combustible and electronic cigarettes. J Am Heart Assoc. 2020, 9(9): e014570

[6] Lee WH, Ong SG, Zhou Y, et al. Modeling cardiovascular risks of e-cigarettes with human-induced pluripotent stem cell-derived endothelial cells. J Am Coll Cardiol. 2019, 73(21): 2722–2737.

[7] Espinoza-derout J, Shao X M, Bankole E, et al. Hepatic DNA damage induced by electronic cigarette exposure is associated with the modulation of NAD+/PARP1/SIRT1 axis. Front Endocrinol (Lausanne). 2019, 10: 320.

[8] Lee HW, Park SH, Weng MW, et al. E-cigarette smoke damages DNA and reduces repair activity in mouse lung, heart, and bladder as well as in human lung and bladder cells. Proc Natl Acad Sci USA. 2018, 115(7): E1560–E1569.

[9] Martin EM, Clapp PW, Rebuli ME, et al. E-cigarette use results in suppression of immune and inflammatory-response genes in nasal epithelial cells similar to cigarette smoke. Am J Physiol Lung Cell Mol Physiol. 2016, 311(1): L135–L144.

[10] Ballbè M, Fu M, Masana G, et al. Passive exposure to electronic cigarette aerosol in pregnancy: A case study of a family. Environ Res. 2022, 8:114490.

[11] Aslaner DM, Alghothani O, Saldaña TA, et al. E-cigarette vapor exposure in utero causes long-term pulmonary effects in offspring. Am J Physiol Lung Cell Mol Physiol. 2022 Oct 11. doi: 10.1152/ajplung.00233.2022. Epub ahead of print. PMID: 36218276.

[12] Ballbè M, Martínez-Sánchez JM, Sureda X, et al. Cigarettes vs. e-cigarettes: Passive exposure at home measured by means of airborne marker and biomarkers. Environ Res. 2014, 135:76–80.

[13] Kyle Strimbu,Jorge A Tavel. What are biomarkers? Current Opinion in HIV and AIDS. 2010, 5(6).

[14] Gerszten Robert E,Wang Thomas J. The search for new cardiovascular biomarkers.Nature. 2008, 451(7181).

[15] Gao, Y., Urine-an untapped goldmine for biomarker discovery? Science China. Life sciences, 2013. 56(12): p. 1145–1146.

[16] Wu J, Li X, Gao Y, et al. Early Detection of Urinary Proteome Biomarkers for Effective Early Treatment of Pulmonary Fibrosis in a Rat Model. Proteomics Clin Appl. 2017, 11(11-12).

[17] Qin W, Li L, Gao Y, et al. Urine Proteome Changes in a TNBS-Induced Colitis Rat Model. Proteomics Clin Appl. 2019, 13(5):e1800100.

[18] Wu J, Zhang J, Gao Y, et al. Urinary biomarker discovery in gliomas using mass spectrometry-based clinical proteomics. Chin Neurosurg J. 2020, 6:11.

[19] Hao Y, Reyes LT, Cheng F, et al. Changes of protein levels in human urine reflect the dysregulation of signaling pathways of chronic kidney disease and its complications. Sci Rep. 2020, 10(1):20743.

[20] Ni M, Zhou J, Zhu Z, et al. A Novel Classifier Based on Urinary Proteomics for Distinguishing Between Benign and Malignant Ovarian Tumors. Front Cell Dev Biol. 2021, 9:712196.

[21] Davies JC, Carlsson E, Midgley A, et al. A panel of urinary proteins predicts active lupus nephritis and response to rituximab treatment. Rheumatology (Oxford). 2021, 60(8):3747–3759.

[22] Meng W, Xu D, Gao Y, et al. Changes in the urinary proteome in rats with regular swimming exercise. Peer J. 2021, 9:e12406

[23] Virreira Winter S, Karayel O, Strauss MT, et al. Urinary proteome profiling for stratifying patients with familial Parkinson’s disease. EMBO Mol Med. 2021, 13(3):e13257.

[24] Virreira Winter, S., et al., Urinary proteome profiling for stratifying patients with familial Parkinson’s disease. EMBO Mol Med. 2021, 13(3): p. e13257.

[25] Watanabe, Y., et al., Urinary Apolipoprotein C3 Is a Potential Biomarker for Alzheimer’s Disease. Dement Geriatr Cogn Dis Extra. 2020, 10(3): p. 94–104.

[26] Huan Y, Wei J, Gao Y, et al. Label-Free Liquid Chromatography-Mass Spectrometry Proteomic Analysis of the Urinary Proteome for Measuring the Escitalopram Treatment Response From Major Depressive Disorder. Front Psychiatry. 2021, 12:700149.

[27] Meng W, Huan Y, Gao Y. Urinary proteome profiling for children with autism using data-independent acquisition proteomics. Transl Pediatr. 2021, 10(7):1765–1778.

[28] Tatemoto K, Hosoya M, Habata Y, et al. Isolation and characterization of a novel endogenous peptide ligand for the human APJ receptor. Biochem Biophys Res Commun. 1998, 251(2):471–476.

[29] Yan J, Wang A, Cao J, et al. Apelin/APJ system: an emerging therapeutic target for respiratory diseases. Cell Mol Life Sci. 2020, 77(15):2919–2930.

[30] Stanisławska-Sachadyn A., Borzyszkowska J., Krzemiński M., et al. Folate/homocysteine metabolism and lung cancer risk among smokers. PLoS ONE. 2019, 14:e0214462.

[31] Giudetti AM, Cagnazzo R. Beneficial effects of n-3 PUFA on chronic airway inflammatory diseases. Prostaglandins Other Lipid Mediat. 2012, 99(3-4):57–67.

[32] Diao WQ, Shen N, Du YP, et al. Fetuin-B (FETUB): a Plasma Biomarker Candidate Related to the Severity of Lung Function in COPD. Sci Rep. 2016, 6:30045.

[33] Aragona P, Puzzolo D, Micali A, et al. Morphological and morphometric analysis on the rabbit conjunctival goblet cells in different hormonal conditions. Exp Eye Res. 1988, 66:81–88

[34] L. Briand, C. Eloit, C. Nespoulous, et al. Evidence of an odorant-binding protein in the human olfactory mucus: location, structural characterization, and odorant-binding properties. Biochemistry. 2002, 41(23):7241–7252

[35] Redl B, Habeler M. The diversity of lipocalin receptors. Biochimie. 2022, 192:22–29.

[36] Huang Y, Jia M, Yang X, et al. Annexin A2: The diversity of pathological effects in tumorigenesis and immune response. Int J Cancer. 2022, 151(4):497–509.

[37] László ZI, Lele Z. Flying under the radar: CDH2 (N-cadherin), an important hub molecule in neurodevelopmental and neurodegenerative diseases. Front Neurosci. 2022, 16:972059.

[38] Lacazette E, Gachon AM, Pitiot G. A novel human odorant-binding protein gene family resulting from genomic duplicons at 9q34: differential expression in the oral and genital spheres. Hum Mol Genet. 2000, 9(2):289–301.

[39] Ezzat Nel-S, Tahoun N. The role of Napsin-A and Desmocollin-3 in classifying poorly differentiating non-small cell lung carcinoma. J Egypt Natl Canc Inst. 2016, 28(1):13–22.

[40] Xi Zhang, QingYu Meng, Jian Jing. Human Annexin A5: from Biomarkers to Therapeutic Candidates. Chinese Journal of Biochemistry and Molecular Biol. 2022, 38(10): 1311–1321.

[41] Nowak JM, Klimaszewska-Wiśniewska A, Izdebska M, et al. Gelsolin is a potential cellular target for cotinine to regulate the migration and apoptosis of A549 and T24 cancer cells. Tissue Cell. 2015, 47(1):105–14.

[42] K. G’al, ’ A. Cseh, B. Szalay, et al. Effect of cigarette smoke and dexamethasone on Hsp72 system of alveolar epithelial cells. Cell Stress Chaperones. 2011, 16:369–378.

[43] A. Hulina-Tomaškovic, I.H. Heijink, M.R. Jonker, et al. Pro-inflammatory effects of extracellular Hsp70 and cigarette smoke in primary airway epithelial cells from COPD patients. Biochimie. 2019, 156:47–58.

[44] M.H. Parseghian, S.T. Hobson, R.A. Richieri. Targeted heat shock protein 72 (HSP72) for pulmonary cytoprotection. Ann. N. Y. Acad. Sci. 2016, 1374:78.

[45] Ponmanickam P, Archunan G. Identification of alpha-2u globulin in the rat preputial gland by MALDI-TOF analysis. Indian J Biochem Biophys. 2006, 43(5):319–22.

[46] M. Yoshida, S. Minagawa, J. Araya, et al. Involvement of cigarette smokeinduced epithelial cell ferroptosis in COPD pathogenesis. Nat. Commun. 2019, 10:1–14.

[47] Eshraghian EA, Al-Delaimy WK. A review of constituents identified in e-cigarette liquids and aerosols. Tob Prev Cessat. 2021, 10;7:10.

